# A genomic and proteomic characterization of mannan-degradable *Bacillus* sp. TTS1, isolated from Tomakomai Forest in Hokkaido

**DOI:** 10.64898/2026.05.18.725066

**Authors:** Saki Mitsumasu, Yu Kasuga, Tatsuya Nagano, Vijay Kumar, Yoshinori Hasegawa, Tomoya Maeda, Taichi E. Takasuka

## Abstract

A challenge in using plant biomass is its highly recalcitrant nature, which makes it economically infeasible to utilize. In natural environments, various microbes, including bacteria and fungi, are reported to decompose plant cell wall materials such as cellulose and hemicellulose, and there may be undescribed microbes that contribute to the degradation of plant biomass. We focused on isolating novel plant biomass-degrading bacteria and screened more than 100 isolates from the Tomakomai experimental forest in Hokkaido, Japan. Among them, one novel *Bacillus* species was chosen for whole-genome sequencing. Comparative genomics and a carbon source utilization assay indicated that the isolate belongs to a subspecies of *Bacillus subtilis*, which we named *B*. sp. TTS1. Glucose, cellobiose, xylose, xylan, mannose, or mannan was used as the sole carbon source in the minimum medium, and the growth of this bacterium was determined. Furthermore, a proteomic analysis of *B*. sp. TTS1 was performed using culture supernatants from various polysaccharide-containing media. In the present study, several key enzymes involved in plant biomass degradation were identified, namely β-1,4-mannanase and xylanase, and they were highly enriched in all tested polysaccharides.

## Introduction

Sustainable production of bio-based fuels and chemicals from plant biomass is needed to address several global challenges, including the high demand for finite petroleum resources, the need to reduce carbon dioxide emissions, and the production of biodegradable chemicals. However, plant cell walls are highly recalcitrant, and the depolymerization of plant biomass into simple sugars still needs high-cost chemical and enzymatic processes for downstream microbial fermentation [1–3]. For example, chemical pretreatment of biomass crops, such as switchgrass, corn stover, and others, needs energy input to operate at high temperatures and pressures and often requires extreme pH conditions [4–6]. Furthermore, dozens of glycoside hydrolases and other enzymes are required for enzymatic depolymerization, thereby challenging the cost-effective saccharification of pretreated plant biomass [2,3,7]. To overcome those technical challenges, it is important to discover novel environmental microbes, determine their plant biomass-degrading abilities, and create mutants to maximize their functions in plant biomass deconstruction processes [7–11]. One of the most successful approaches was applied to *Trichoderma reesei* QM6a (syn. *Hypocrea jecorina*), isolated from Solomon Islands during World War II [12–14], which hydrolyzes not only cellulose and hemicellulose in the plant cell wall but also depolymerizes lignins. Tremendous efforts were made to develop a recent industrial strain, *T. reesei* RutC-30 and other strains, which enable the production of plant biomass-degrading enzymes in high quantities, and the enzymatic cocktails are commercially available [15–17]. However, cost-effective plant biomass depolymerization has not been achieved to date, and it remains important to discover novel microbes from Nature that meet the demand for efficient enzymatic degradation of plant biomass [18,19].

In the present study, we attempted to isolate plant biomass-degrading bacteria from the Hokkaido forest and identified a novel bacterium, *B*. sp. TTS1. Phylogenetically, the strain was closely related to the *Bacillus subtilis* type strain. Consistently, carbon utilization showed overall similar, but a few differences. *B*. sp. TTS1 grew well on a series of different carbon sources, and secretomic analysis determined key enzymes that might hydrolyze polysaccharides in the plant cell walls. Overall, the present study indicates possible roles of *B*. sp. TTS1 as an environmental bacterium that utilize the plant cell wall components, especially mannan and xylan.

## Result and Discussion

### Novel bacterial isolates from Tomakomai Forest in Hokkaido

We isolated over a hundred microbes from the Hokkaido University Experimental Forest in Tomakomai City, Japan (**Fig. 1a**), and several isolates grew on a minimal medium containing carboxymethyl cellulose (CMC) as the sole carbon source. We chose one isolate and performed 16S rRNA sequencing, which identified it as *Bacillus* sp., hereafter referred to as *B*. sp. TTS1. *B*. sp. TTS1 grew well on glucose, locust bean gum (LBG), or xylan as a sole carbon source, but less so on CMC-containing M9 minimum medium (**Fig. 1b**).

**Fig. 1.**
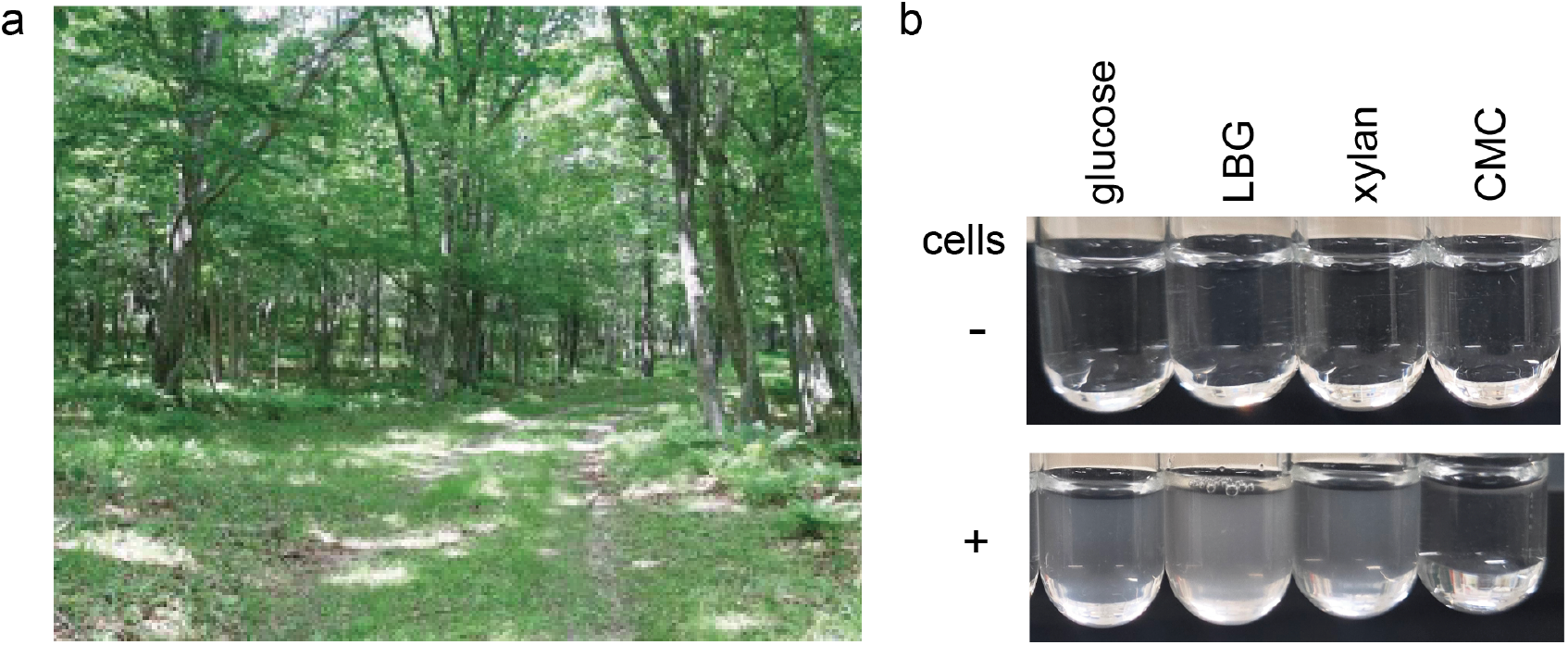
Novel isolates obtained from Tomakomai Forest and their ability to utilize selected carbon sources. **a**. *B*. sp. TTS1 was isolated from Tomakomai Forest in Hokkaido. **b**. *B*. sp. TTS1 was grown on M63 minimum medium containing either glucose, LBG, xylan, or CMC as a sole carbon source.

A whole-genome sequencing of *B*. sp. TTS1

The whole genome of *B*. sp. TTS1 was sequenced. The 4,211,021 bp genome was *de novo* assembled, and the overall G+C content was estimated to be about 43.6 % (Table 1). The gene annotation resulted in 4,176 protein-coding genes, of which 134 (3.2% of the total proteome) were predicted to be vCarbohydrate-Active Enzymes (CAZymes) [20,21], including 54 Glycoside Hydrolases (GHs), 7 Polysaccharide Lyases (PLs), 18 Carbohydrate Esterases (CEs), and 11 Carbohydrate Binding Modules (CBMs). *B*. sp. TTS1, with five *Bacillus* species, including *B. subtilis* NBRC 13719, *B. amyloquefaciens* NBRC 15535, *B. pumils* NBRC 12092, and *B. thuringiensis* NBRC101235, shows that the genome of *B*. sp. TTS1 is closest to the *B. subtilis* NBRC 13719 strain, with a whole-genome average nucleotide identity (ANI) value of 98.8 and a digital DNA-DNA hybridization (dDDH) value of 89.9 (Table 1). Therefore, it was concluded that the *B*. sp. TTS1 belongs to a member of the *B. subtilis* species with a few distinctive features, including the number of proteomes, GHs, and CBMs. For instance, the number of protein-coding genes, GHs, and CBMs encoded in the *B*. sp. TTS1 genome was 133, 4, and 3 fewer than in the genome of *B. subtilis* NBRC 13719, respectively. As reported elsewhere on the diversity of *Bacillus* species, the genome contents of other *Bacillus* species differ substantially from those of *B*. sp. TTS1 and *B. subtilis* NBRC 13719 (Table 1). Several important strains have been commercialized, such as *B. subtilis* var. natto an important bacterium for fermented soybeans, which is used to produce natto and natto kinase with strong fibrinolytic activity [22], *B. thuringiensis* for producing biopesticides, and other *Bacillus* species. for bioplastic productions [23]. Furthermore, some *Bacillus* species were reported to utilize different types of sugars and polysaccharides, including, cellulose, mannan, xylan, keratin, polythene, and others [23–25]. Therefore, the following phylogenetic, proteomic, and biochemical analyses were performed to characterize *B*. sp. TTS1 in more detail.

**Table 1.**
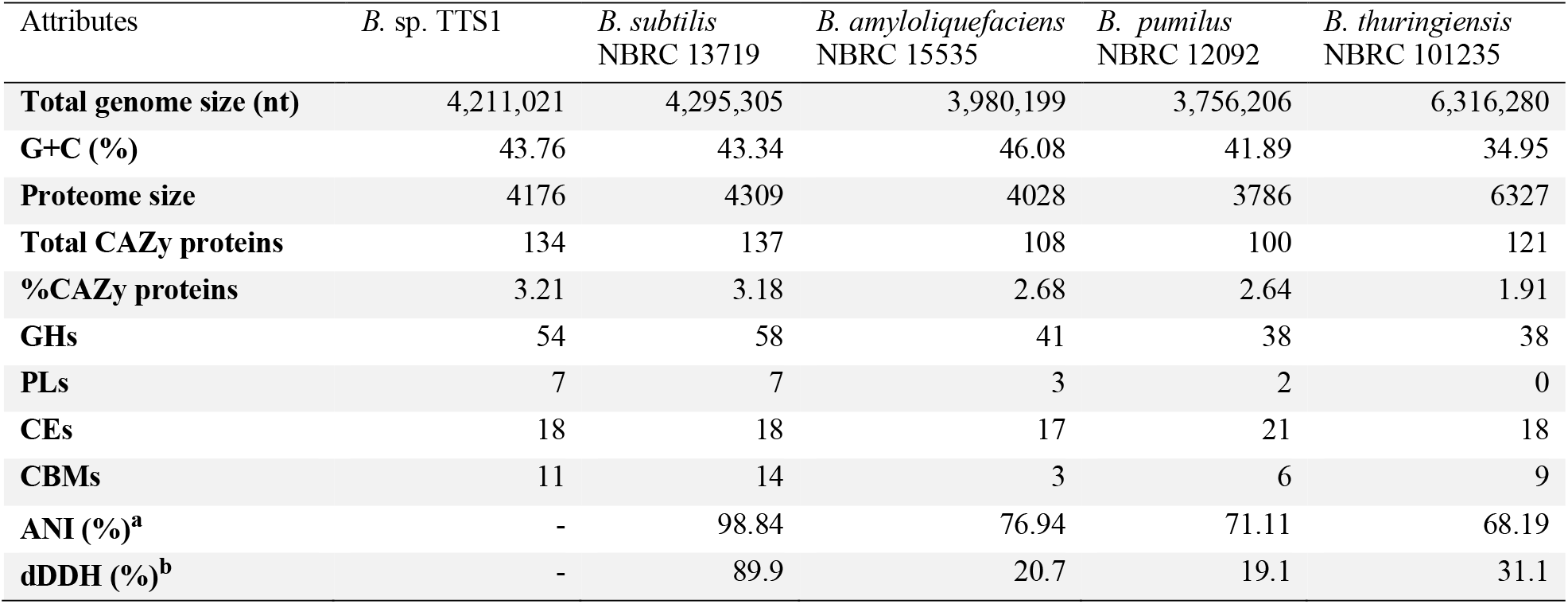
Genome analysis of *B*. sp. TTS1 and comparison with other *Bacillus* spp. *B*. sp. TTS1 and four other type strains are shown with total genome size, G+C (%), proteome size, CAZyme potential, including total number of CAZymes, %CAZymes-coding genes relative to total proteome numbers, number of GHs, PLs, CEs, and CBMs, respectively. ^a^ANI (%) and ^b^dDDH indicate the values estimated in each strain relative to *B*. sp. TTS1.

### Phylogenetic analysis of *B*. sp. TTS1 and other *Bacillus* type strains

To analyze the phylogenetic distance of *B*. sp. TTS1 to other *Bacillus* spp., a 16s rRNA-based phylogenetic tree was constructed (**Fig. 2a**). It showed that *B*. sp. TTS1 was close to *B. tequilensis* NBRC 101235, and *B. subtilis* NBRC 13719. The similarity of 16S rRNA among these 3 strains was significantly high, so a more detailed analysis was conducted using the whole-genome sequence. As shown in Table 1, ANI and dDDH values indicated that the *B*. sp. TTS1 belongs to a novel subspecies of *B. subtilis*. Pairwise comparison of genome sequences, phylogenetic inference, and type-based species and subspecies clustering across *B*. sp. TTS1 and other *Bacillus* strains were performed, respectively (**Fig. 2b**). The results also support that *B*. sp. TTS1 is a *B. subtilis* subspecies with a small phylogenetic distance.

**Fig. 2.**
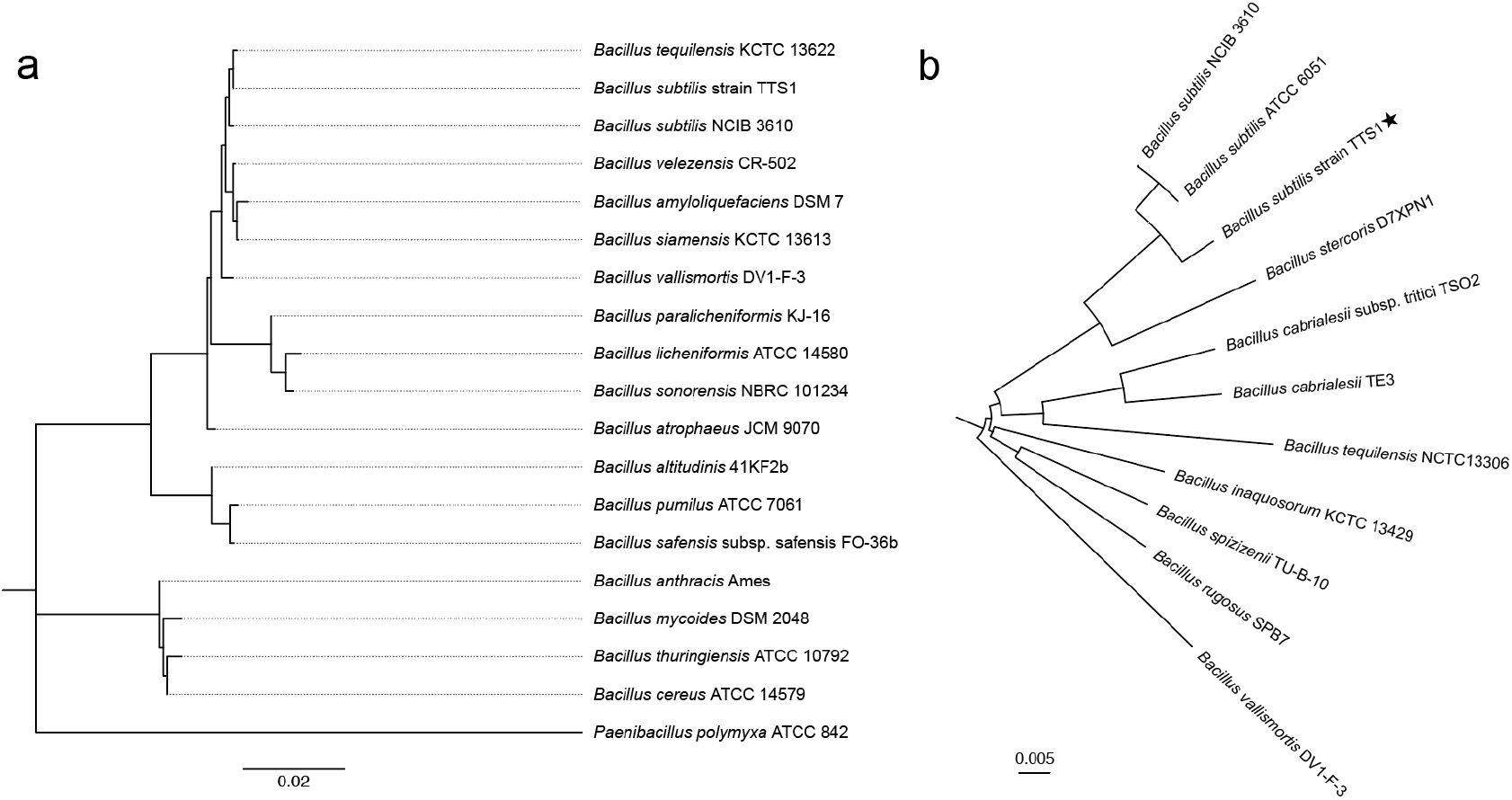
Phylogenetic trees showing relationships of *B*. sp. TTS1 and other *Bacillus* type strains. **a**. The phylogeny was constructed from 16S rDNA sequences of *Bacillus* species and inferred using the maximum-likelihood method (bootstrap=1,000). **b**. Multilocus alignment phylogeny inferred from GBDP distances with GBDP pseudo-bootstrap support values > 60% from 100 replications, with an average branch support of 72.4%, and the tree was rooted at the midpoint.

### Sugar utilization of *B*. sp. TTS1 with other *Bacillus* spp

Further characterization of *B*. sp. TTS1 was performed using the API® 50 CH, which enables the determination of bacterial organic acid production in response to a series of different carbon sources along with 4 *Bacillus* type strains, including *B. subtilis* NBRC 13719, *B. amyloquefaciens* NBRC 15535, *B. pumils* NBRC 12092, and *B. thuringiensis* NBRC 101235 (Table 2). Each *Bacillus* strain showed different outcomes across a series of carbon sources and especially for phylogenetically distant strains, including *B. amyloquefaciens* NBRC 15535, *B. pumils* NBRC 12092, and *B. thuringiensis* NBRC 101235 from *B*. sp. TTS1. Between *B*. sp. TTS1 and *B. subtilis* NBRC 13719, the phenotypic pattern against most of the tested sugars was similar, except for gentiobiose, which was utilized only by *B. subtilis* NBRC 13719, potentially differentiating the two *Bacillus* species.

**Table 2.**
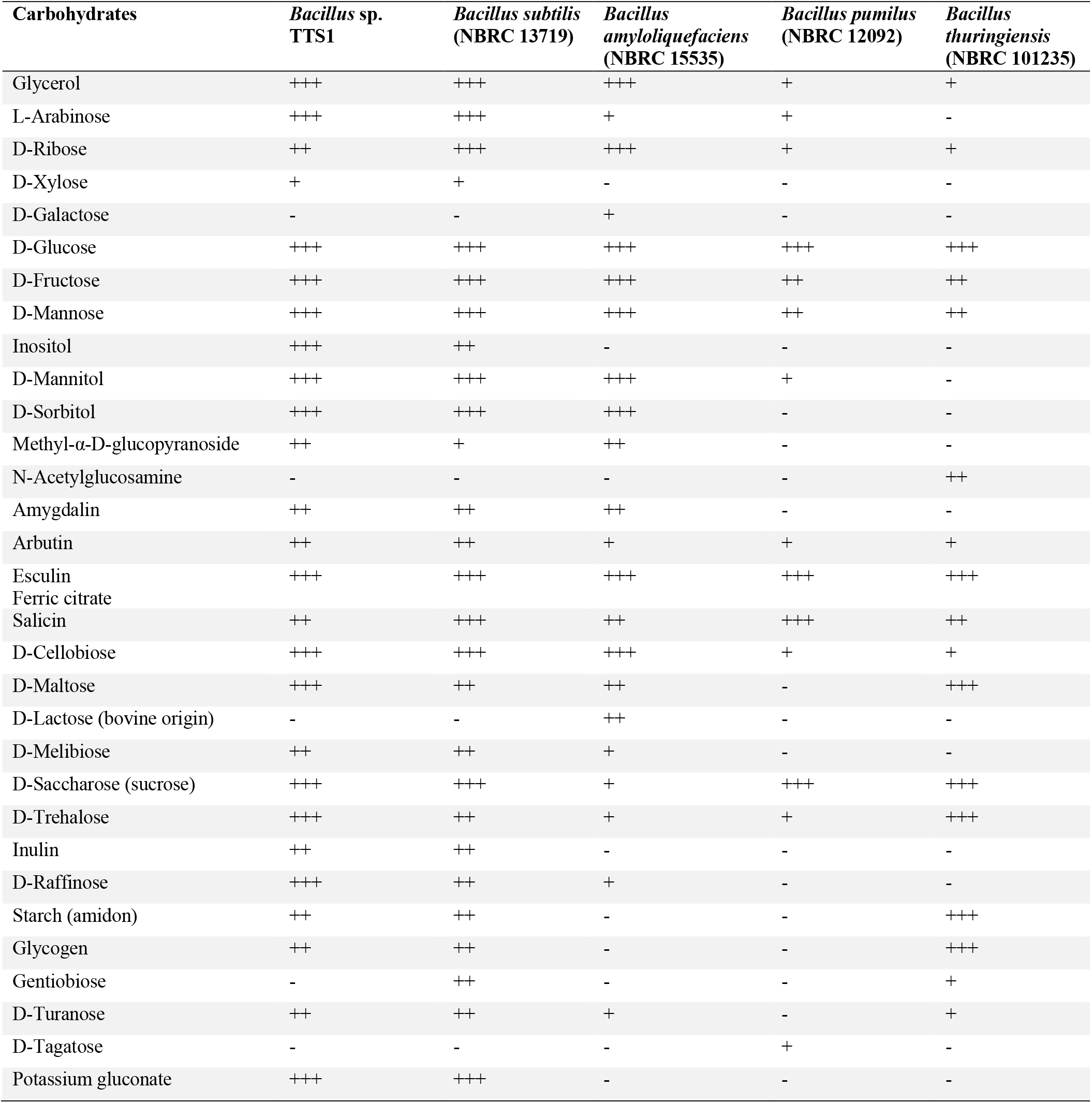
The difference in carbohydrate metabolism between *B*. sp. TTS1 and other *Bacillus* spp. was observed by biochemical assay. The degree of medium color changes corresponding to the production of organic acids from carbohydrate consumption is shown as +/-. “+++”, “++”, and “+” indicate the degree of growth and “-” indicates the lack of utilization of the corresponding carbohydrate.

### Measurements of the cell growth of *B*. sp. TTS1 in the presence of plant cell wall constituents

To determine the growth of *B*. sp. TTS1 in the presence of plant cell wall materials, including glucose, cellobiose, mannose, LBG, xylose and xylan, cells were grown on M63 minimum media with indicated carbon sources for every 10 min for 3 days along with *B. subtilis* NBRC 13719 (**Fig. 3, Supplementary data 1**). To note, the culture containing CMC as the sole carbon source showed very little growth of both strains, and *B*. sp. TTS1 and *B. subtilis* NBRC 13719 showed similar growth trends (**Supplementary data 1**). In the presence of glucose or cellobiose, the cells reached a plateau at around 30 h with 0.4OD_600_ and 24 h with 0.5OD_600_, respectively, in *B*. sp. TTS1. In contrast, when cells were grown on xylose, they reached a plateau significantly more slowly, with an OD_600_ of less than 0.2, even compared to growth on xylan, where cells reached a plateau at around 14 h with an OD_600_ of 0.3. Interestingly, when mannose was used as the sole carbon source in the medium, growth appeared to enter the stationary phase at around 14 h with an OD_600_of 0.3, paused between 15 and 21 h, then regrew until the end point with an OD_600_of 0.4 (72 h). This observation might reflect the activation of the reported mannose-utilizing operon in several *B. subtilis* strains, which include the mannose PTS transporter, mannose-6-phosphate isomerase, and mannose transcriptional regulator for adaptation to mannose [26–28]. Thus, it is thought that the transporter uptake and phosphorylates mannose, followed by the isomerization of mannose-6-phosphate into fructose-6-phosphate for glycolysis. Interestingly, cell growth on LBG differed clearly from that on other carbon sources used in the media. A significant spike in OD_600_ was observed at around 20 h, reaching ∼1.5. This observation was thought to be due to cell-to-LBG aggregation. The optical density decreased to ∼0.8 between 20 and 24 h, and cells appeared to enter the stationary phase, which might be explained by the enzymatic hydrolysis of LBG during growth. *B. subtilis* NBRC 13719 showed slightly more growth on glucose, cellobiose, xylose, and mannose. Interestingly, neither *B*. sp. TTS1 nor *B. subtilis* NBRC 13719 utilized galactose (Table 2), one of the sugars in LBG, hence, cell growth should be supported only by mannose, yet it is not clear whether they utilize galactosyl-mannose or other oligosaccharides, such as mannobiose, mannotriose, galactosyl-mannobiose, and others.

**Fig. 3.**
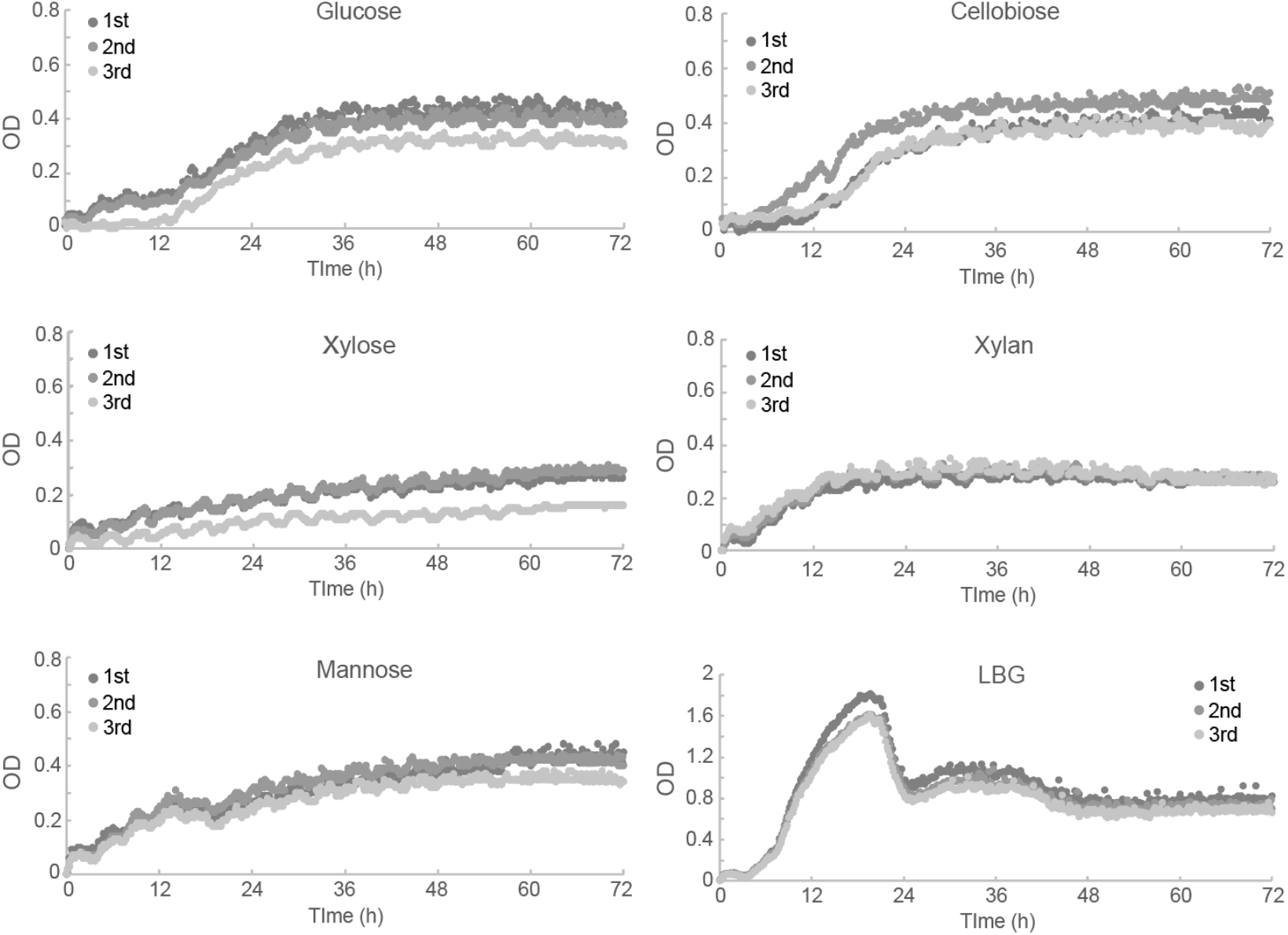
Growth assay of *B*. sp. TTS1 in the presence of different carbon sources. Cells were grown on M63 minimum medium with the indicated sole carbon source, including glucose, cellobiose, xylose, xylan, mannose, or LBG. Three independent experiments are shown as 1st, 2nd, and 3rd plots, respectively.

### Proteomic analysis of the *B*. sp. TTS1 cells grown on different carbon sources

A liquid chromatography-tandem mass spectrometry (LC/MS-MS) analysis for *B*. sp. TTS1 grown in the following carbon sources, including glucose, cellobiose, sigmacell cellulose, beechwood xylan, and purified mannan, was performed. Notably, *B*. sp. TTS1 grew and produced detectable amounts of proteins when grown on Sigmacell cellulose as the sole carbon source, but not on CMC. Overall, 1041, 1142, 630, 1013, and 1008 proteins were determined in glucose, cellobiose, sigmacell cellulose, beechwood xylan, and purified mannan samples, respectively (Table 3). Among those proteins, 54, 56, 45, 64, and 68 possess signal peptide sequences, and the rest were considered to be intracellular proteins in glucose, cellobiose, sigmacell cellulose, beechwood xylan, and purified mannan samples, respectively. The top 18 predicted extracellular CAZymes with a predicted signal peptide were extracted and sorted in descending order from the mannan dataset (**Fig. 4**). The exponentially modified protein abundance index (emPAI) was used to represent protein abundance for each sample across three independent analyses [29]. BTTS1_3214 was the most enriched protein with annotated β-1,4-mannanase belonging to the GH26 family, followed by GH11 xylanase (BTTS_1149) in all carbon sources. Other 16 secreted CAZymes include two GH43 α-L-arabinosidases (BTTS_0217 and BTTS_2122), one each of GH46 chitosanase (BTTS1_0484), GH30_8 xylanase (BTTS_4004), GH13_28 α-amylase associated with CBM26 (BTTS_3502), PL3_1 pectate lyase (BTTS_1664), CBM63 expansin (BTTS_1173), GH18 chitosanase (BTTS_1571), GH16_21 lichenase (BTTS_2094), PL1_6 pectate lyase (BTTS_3757), GH53 β-1,4-galactanase (BTTS_1581), GH5_2 endoglucanase associated with CBM3 (BTTS_4002), GH171 β-N-acetylmuramidase (BTTS_3636), GH32 levanase associated with CBM66 (BTTS_0469), PL1_8 pectate lyase (BTTS_1171), and GH73 β-N-acetylglucosaminidase (BTTS_1757).

**Table 3.**
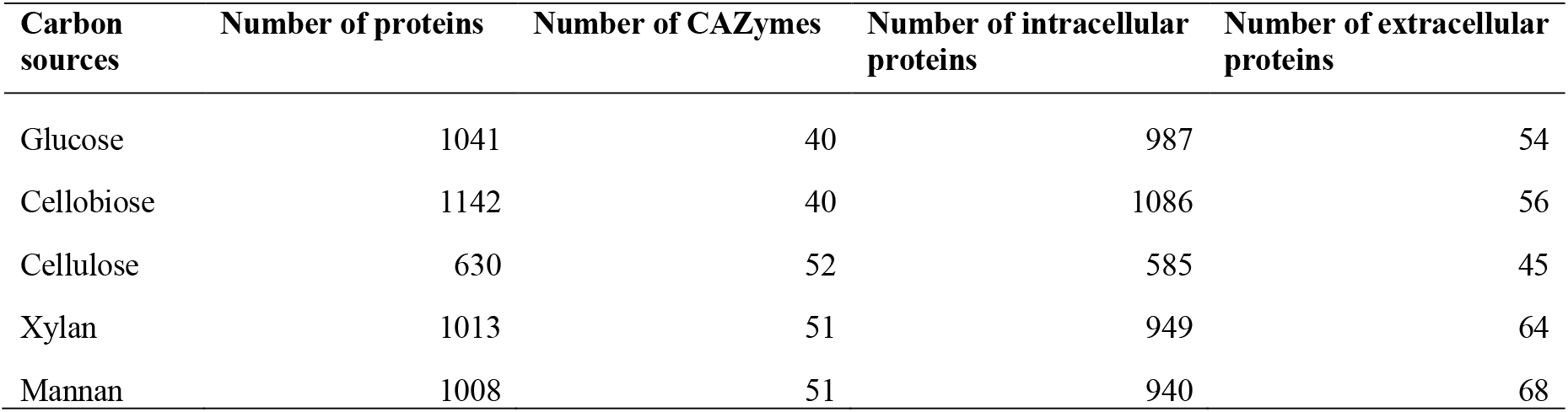
Summary of proteomics analysis of *B*. sp. TTS1 grown on different carbon sources. The number of predicted total proteins, extracellular proteins, intracellular proteins, and predicted CAZymes is shown.

**Fig. 4.**
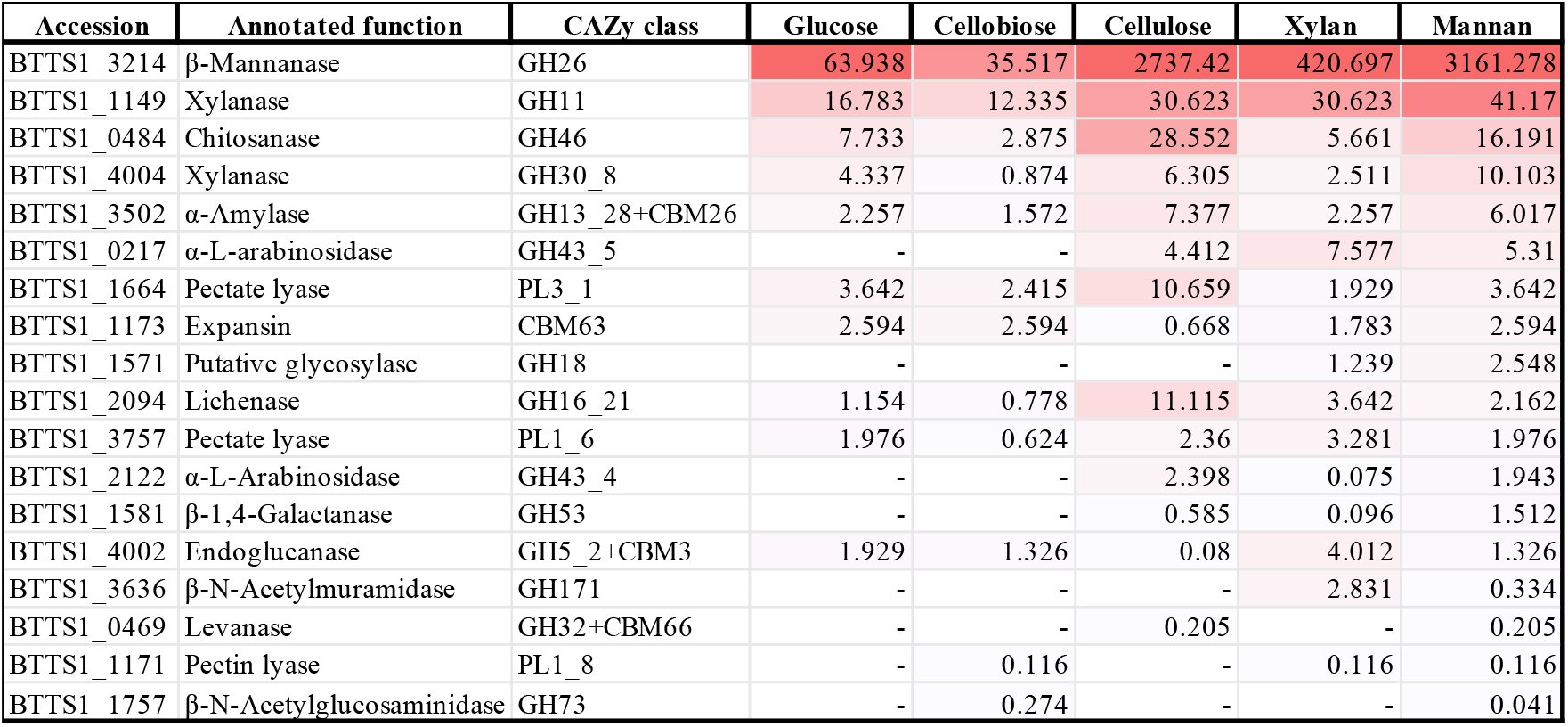
Secretomic analysis for TTS1 revealed that β-mannanase was secreted on cellulose and xylan medium as well as mannan medium. Annotated protein functions and CAZyme classifications for each protein are shown, along with calculated emPAI values from three biological replicates, and the dataset is sorted by mannan culture supernatant.

Although most of the predicted extracellular CAZymes were detected in the glucose- and cellobiose-grown cells, the protein abundance based on the emPAI was relatively lower than other datasets (**Fig. 4**). Therefore, it is concluded that *B*. sp. TTS1 produces a limited amount of extracellular CAZymes in response to simple sugars, consistent with the previous reports [30,31]. An extremely high abundance of β-1,4-mannanase (BTTS_3214), a homolog of *B. subtilis* 168, gmuG, was determined in cellulose-, xylan-, and mannan-grown cells. Consistently, gmuG was detected in an earlier study under cellobiose-grown conditions of *B. subtilis* 168 [31]. In polysaccharide-grown culture supernatants, GH11 xylanase (BTTS_1149) and GH46 chitosanase (BTTS1_0484) were the second and third most abundant proteins, as reported in previous studies [30,31]. In the cellulose culture supernatant, two CAZymes, PL3_1 pectate lyase (BTTS_1664) and GH16_21 lichenase (BTTS_2094) were secreted with the emPAI values > 10. In the mannan culture supernatant, other predicted secreted CAZymes were detected, but at lower levels compared to β-1,4-mannanase (BTTS_3214). The transcriptional regulation of the gene cluster, containing the homolog of this enzyme in *B. subtilis* 168, gmuG, was reported, and a helix-turn-helix type transcriptional repressor, gmuR, was reported to be released from the promoter region in response to cellobiose, mannobiose or other glucomannan-derived oligosaccharides [32]. The current result also indicates that xylooligosaccharides and other oligosaccharides in beechwood xylan might also induce the expression of downstream genes by gmuR [33]. Our results also suggest that longer oligosaccharides, such as -triose, tetraose, and others, are more effective ligands to derepress the β-1,4-mannanase (BTTS_3214) gene, since the protein abundance in the cellobiose culture supernatant was significantly lower compared to that of polysaccharides. The secretomic results indicated that the detected β-1,4-mannanase (BTTS_3214) might function in the hydrolysis of multiple polysaccharides, such as cellulose and mannan.

## Conclusions

In this study, we isolated a novel bacterium from the forest in Hokkaido, Japan. Based on the 16S rDNA sequencing, while genome sequencing, and sugar utilization assay, the isolate was determined to be a subspecies of *B. subtilis*, and named *B*. sp. TTS1. The growth test of *B*. sp. TTS1 supports that this bacterium grows well on glucose, cellobiose, xylose, xylan, mannose, and LBG to different degrees. The secretomic analyses identified a potential key enzyme for depolymerizing polysaccharides, including cellulose, xylan, and mannan. Further study is needed to determine whether the detected β-1,4-mannanase (BTTS_3214) might possess multifunction to hydrolyze different polysaccharides.

## Materials and Methods

### Reagents

A D-glucose (Nacalai Tesque, Kyoto, Japan), D-cellobiose (Biosynth, Compton, UK), D-Xylose (Tokyo Chemical Industry, Tokyo, Japan), and Sigmacell Cellulose (Sigma-Aldrich, St. Louis, MO) were purchased. A D-mannose, D-galactose, carboxymethyl cellulose (CMC), and locust bean gum were purchased from Wako Pure Chemicals (Osaka, Japan). Beechwood xylan and D-mannan were purchased from Megazyme (Bray, Ireland).

### Bacterial isolation

Environmental bacteria from Tomakomai Forest were screened on an M63 minimum medium agar plate containing 1% CMC as the sole carbon source for 3 days at 30 °C. Cells were streaked multiple times on a 1% CMC M63 plate until isolates were obtained and visually confirmed under an optical microscope.

### DNA extraction

Bacterial DNA from *B*. sp. TTS1 was extracted from pure culture using phenol chloroform extraction method. *B*. sp. TTS1 was cultured in 20 mL YME medium containing 4 g yeast extract, 10 g malt extract, and 4 g D-glucose in 1 L distilled water, for 2 days at 30°C. Cells were harvested by centrifugation at 4,000 x *g* for 10 min and washed once with 10% sucrose buffer and suspended in 250 mL of Lysis solution (25 mM EDTA, 25 mM PH 8.0 Tris-HCl buffer, and 0.3 M sucrose), 12 U/mL RNase A (Macherey-Nagel GmbH & Co.,, Mannheim, Germany), and 1 mg/mL lysozyme (Wako Pure Chemicals, Osaka, Japan) and incubated for 40 min at 37 °C. 10% SDS and 10 mg/mL proteinase-K (FIJIFILM Wako Pure Chemical Corporation, Osaka, Japan) were added and inverted several times, followed by incubation for 2 h at 55 °C. An equal amount of phenol chloroform solution (phenol:chloroform:isoamyl alcohol/25:24:1) was added and gently mixed. After centrifugation for 10 min at 15,000 x *g*, the upper layer was collected, and the process was repeated until the white protein layer disappeared. The extracted DNA was ethanol precipitated and dissolved in TE buffer. The purity and amount of extracted DNA were quantified by using a spectrophotometer (DeNovix DS-11, Wilmington, DE).

### Whole genome sequencing and gene annotation of *B*. sp. TTS1

A DNA library was prepared from 1 μg of genomic DNA using MGI Easy PCR-Free Library Prep Set (MGI, China). Sequencing was performed on a DNBSEQ-G400 (MGI), using paired-end 150 bp reads and de novo assembly was performed. CDSs were extracted using Prokka [34], and their functions were annotated by BLAST sequence similarity searches. The secreted proteins were predicted by SignalP-6.0 [35], and the CAZyme functions were annotated by dbCAN [20,21,36].

### Construction of phylogenetic trees

16S rRNA sequences of *Bacillus* type strains were obtained from Ezbiocloud [37] and phylogenetic analyses were conducted using MEGA version 12.1 [38]. For a whole genome-based taxonomic analysis, the genome sequence data were uploaded to the type strain Genome Server (TYGS, https://tygs.dsmz.de) [39]. Pairwise comparison of genome sequences [40,41], phylogenetic inference [42], and type-based species and subspecies clustering [43] between *B*. sp. TTS1 and other *Bacillus* strains were performed, respectively.

### Phenotypic assay

The bacterial strains, *B. subtilis* NBRC 13719, *B. amyloliquefaciens* NBRC 15535, *B. pumilus* NBRC 12092, and *B. thuringiensis* NBRC 101235, were obtained from the National Biological Resource Center (NBRC) (NITE, Tokyo, Japan). Aerobic acid production due to carbohydrate utilization was tested using API® 50 CH with API® 50 CHB/E Medium (bioMérieux, France) according to the manufacturer’s instructions. Cells were grown in 2 mL of YME medium for 2 days at 30 °C and harvested by centrifugation for 10 min at 5,000 x *g*. Cells were adjusted to 0.5 OD/mL in 10 mL of API® 50 CHB/E Medium, inoculated into 50 reaction pockets, and incubated for 24 h at 30 °C.

### Measurement of cell growth

Preculture of *B*. sp. TTS1 was prepared in YME medium for overnight at 30 °C, and cells were collected by centrifugation for 10 min at 4,000 x *g*. The cells were washed once with the M63 minimal medium (10.72 g dipotassium hydrogen phosphate, 5.24 g potassium dihydrogen phosphate, 2 g ammonium sulfate in 1 L of distilled water) and adjusted to 0.1 OD_600_. The cells were inoculated into the M63 minimal medium supplemented with 1 mM magnesium sulfate, 1 mg thiamin, and one of the following carbon sources in 0.5% final concentration: glucose, cellobiose, mannose, LBG, xylose or beechwood xylan. Cell growth was monitored using a spectrophotometer Epoch2 (Agilent, Santa Clara, CA) and plotted every 10 min.

### Preparation of extracellular proteomic analysis samples

*B*. sp. TTS1 was cultured for 3 days at 30 °C in M63 minimal medium supplemented with 1% each of glucose, cellobiose, sigmacell cellulose, beechwood xylan, or mannan. The culture supernatants were collected by centrifugation at 5000 x *g* for 10 min at 4 °C. Secreted proteins in each culture supernatant were precipitated by adding 25 % (vol/vol) trichloroacetic acid (TCA), washed twice with ice-cold acetone, and denatured with 8 M urea. 25 mM ammonium bicarbonate was added to 20 μg of proteins to dilute into 1 M urea, and proteins were reduced with 50 mM dithiothreitol for 30 min at 50 °C and alkylated with 500 mM iodoacetamide for 30min in the dark at room temperature. Trypsin (proteomic grade; Roche, Mannheim, Germany) was added (1:40 trypsin/secretome ratio) and incubated for 16 h at 37 °C and purified with C_18_ ZipTip pipette tips (Millipore, Ireland).

### Proteomic analysis

Proteomic analysis was performed as described previously [44–46]. Briefly, mass spectra of extracellular proteins were obtained by an EAZY-nLC 1000 liquid chromatography system with Q Exactive Plus Orbitrap mass spectrometer (Thermo Fisher Scientific, Illinois) and Xcalibur software v. 3.0 (Thermo Fisher Scientific). The peptides were separated on a C_18_ capillary tip column (NTCC-360/75-3-125; Nikkyo Techno, Japan) by linear gradient from 5 to 30% with two solutions, 0.1% formic acid in water (solution A) and 0.1% formic acid in acetonitrile (solution B), for 120 min. Full-scan mass spectra were acquired in the Orbitrap mass spectrometer with a scan range of 300.0 to 2,000.0 m/z with a resolution of 70,000. Proteins were identified from the MS/MS spectra using Proteome Discoverer v. 2.1.0.81 (Thermo Fisher Scientific) with the CDSs of *B*. sp. TTS1 (**Supplementary data 2**), and the exponentially modified protein abundance index (emPAI) was estimated [29,44,45,47]. The peptide mass tolerance was set at 10 ppm, and the fragment mass tolerance was set at 0.8 Dalton, respectively. The peptide charge was set at 12, 13, and 14. The accuracy and sensitivity of peptide identification were optimized using the automatic decoy and percolator functions built into the Proteome Discoverer software. All proteomics data sets are accessible via the PRIDE proteomics identification database.

## Acknowledgements

This work was supported by the JSPS KAKENHI Grant Number 23K27036, 25K22375, and JST ALCA-Next (FS) Grant Number JPMJAN24D6 (T.E.T.), Grant-in-Aid for JSPS Research Fellow Grant Number JP 24KJ0307 (T.N.) and JSPS Postdoctoral Fellowships for Foreign Researchers (Standard Program) 20F20391 P20391 (V.K.).

## Author contributions

S.M., V.K., T.M., and T.E.T. designed the experiments. S.M., Y.K., T.N., and Y.H. performed experiments and analyzed the results. S.M., Y.K., T.N., Y.H. V.K., T.M., and T.E.T. wrote and edited the manuscript. All the authors agreed on the manuscript.

## Conflict of interest

The authors declare no conflict of interest.

## Data Availability

All proteomic data sets obtained in this study can be accessed via the PRIDE proteomics identification database (https://www.ebi.ac.uk/pride/) under the accession number PXDXXXXXX for proteomics analysis.

## Notes

### Competing Interest Statement

The authors have declared no competing interest.

